# Diaphragmatic EMG Uncovers Left-Right Diaphragmatic Imbalance After Ischemic Stroke with Implications for Motor Recovery

**DOI:** 10.64898/2026.07.10.737872

**Authors:** Ziyi Feng, Jieqiong Wang, Qianzi Cong, Junhui Bai, Yuehua Zhao, Jie Zhu, Qingming Qu, Jie Jia

## Abstract

**Background:** Stroke-associated respiratory dysfunction has been increasingly recognized, yet whether diaphragmatic impairment after stroke involves lateralized neuromuscular imbalance remains unclear.

**Methods:** In this controlled experimental study, transient left middle cerebral artery occlusion and reperfusion (tMCAO) was performed in mice, followed by multimodal assessment of treadmill-based peak oxygen uptake testing, diaphragm ultrasonography, bilateral diaphragmatic electromyography (dEMG) with simultaneous respiratory flow monitoring, behavioral assessment, and whole-mount immunofluorescence imaging of the diaphragm.

**Results:** Ischemic stroke reduced exercise capacity and diaphragmatic excursion without detectable diaphragm thinning, indicating functional impairment rather than overt atrophy. Bilateral dEMG revealed a subacute left–right asymmetry in inspiratory activation, with the ipsilesional hemidiaphragm exhibiting reduced amplitude, decreased area under the curve, and altered burst duration, consistent with a relative contralateral-dominant pattern rather than frank hyperactivation. Whole-mount imaging demonstrated asymmetric nerve remodeling, characterized by a nadir in ipsilesional nerve fiber density at 1 week and partial recovery by 2 weeks after stroke. These structural changes closely paralleled the dEMG alterations, suggesting a neural substrate for lateralized dysfunction. Exploratory analyses further linked dEMG parameters with motor recovery.

**Conclusions:** These findings provide novel evidence that ischemic stroke induces a left–right diaphragmatic neuromuscular imbalance, with structural and functional correlates in diaphragmatic innervation. The study further supports the utility of bilateral dEMG as a functional marker for assessing diaphragmatic dysfunction and monitoring recovery after stroke.

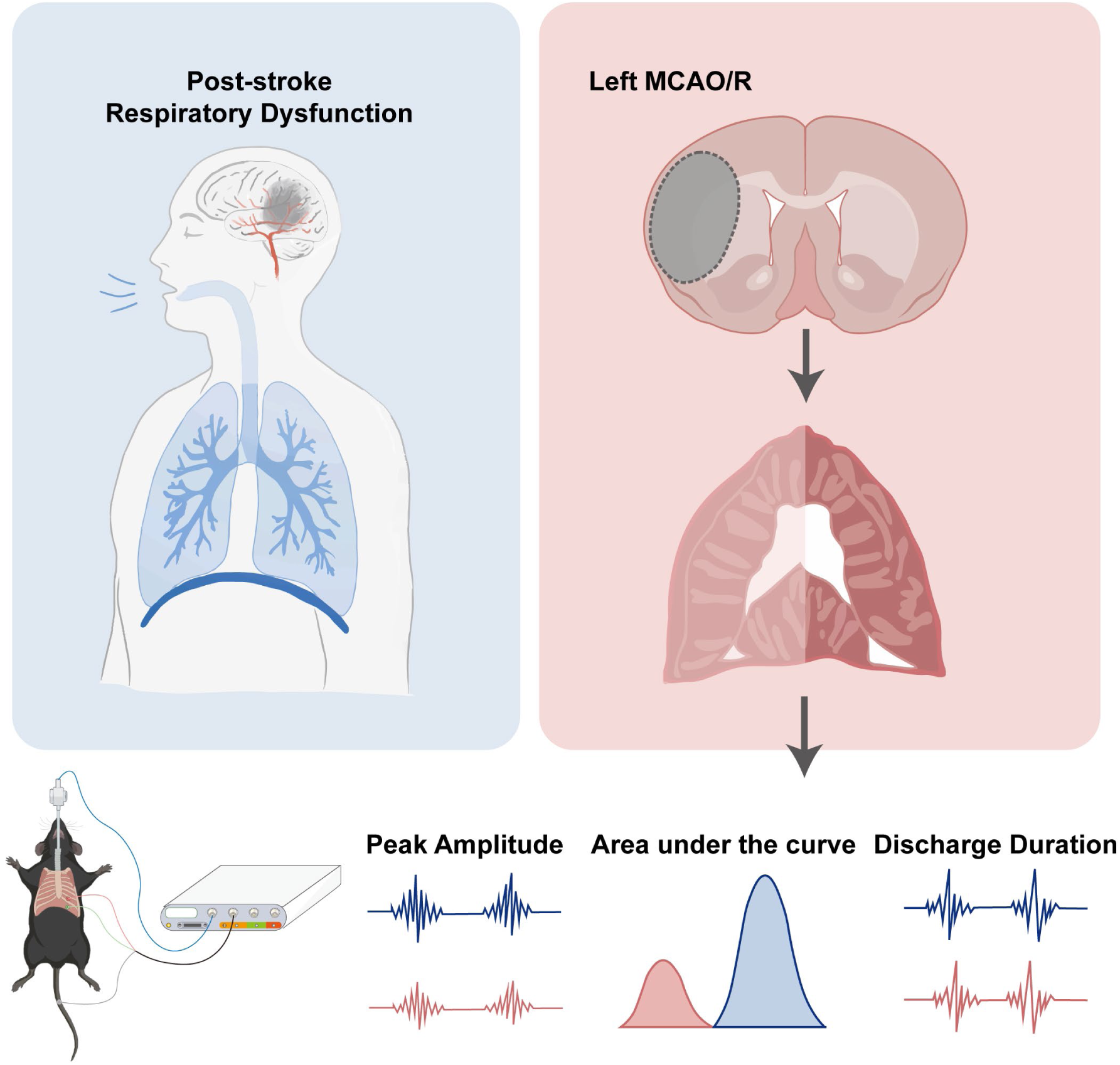

## Introduction

Stroke remains a leading cause of long-term disability and mortality worldwide^1^. Although post-stroke impairments are commonly characterized by motor, sensory, language, and cognitive impairments, respiratory dysfunction has been increasingly recognized as an important yet underappreciated complication^2^. Stroke patients exhibit reduced pulmonary function, impaired respiratory muscle strength, diminished cough efficiency, ventilatory inefficiency, and decreased exercise tolerance^3^. These abnormalities can increase the risk of pulmonary complications, restrict participation in rehabilitation, and ultimately contribute to poor functional outcomes^4–9^.

Current clinical and experimental assessments, including cardiopulmonary exercise testing, pulmonary function testing, and respiratory muscle strength assessment, are useful for detecting global respiratory impairment after stroke^10^. However, these approaches provide limited insight into the specific neuromuscular mechanisms underlying respiratory dysfunction^11^. Breathing is regulated by an integrated motor system involving central respiratory networks, supraspinal descending pathways, spinal motor neurons, phrenic nerve output, respiratory muscles, thoracic mechanics, and pulmonary–cardiovascular interactions^12^. Consequently, the pathway through which focal cerebral injury disrupts respiratory performance remains incompletely understood. In particular, whether unilateral stroke alters the bilateral coordination of respiratory motor output has not been fully clarified^11,13^.

The diaphragm is the principal inspiratory muscle and plays a central role in maintaining effective ventilation. Normal diaphragmatic contraction depends on coordinated bilateral activation through the phrenic nerves, which are regulated by brainstem respiratory networks and influenced supraspinal descending pathways^14–16^. Given that stroke often involves unilateral cerebral injury, disruption of descending motor control may disturb the balance of bilateral diaphragmatic neural drive^17^. Emerging clinical evidence supports this possibility. Studies using surface electromyography and ultrasonography have demonstrated asymmetric diaphragmatic activation in patients with stroke, with the degree and pattern of asymmetry varying under different inspiratory loads^18,19^. Such lateralized diaphragmatic dysfunction may compromise ventilatory efficiency and represent a previously underrecognized mechanism of post-stroke respiratory impairment. Beyond its contribution to respiratory dysfunction, diaphragmatic neuromuscular imbalance may also have relevance to neurological recovery. Because diaphragmatic activation depends on the integrity of central–peripheral motor pathways, altered bilateral diaphragmatic drive may reflect broader stroke-induced disruption of motor network function^20^. Accordingly, diaphragmatic electrophysiological parameters may provide physiological information not only about respiratory impairment but also about recovery-related neuromuscular status^21^. However, direct evidence linking focal cerebral ischemia to bilateral diaphragmatic neural drive imbalance, structural remodeling of diaphragmatic innervation, and functional recovery remains limited.

Therefore, the present study investigated bilateral diaphragmatic function after focal cerebral ischemia using a multimodal approach combining exercise testing, diaphragm ultrasonography, bilateral diaphragmatic electromyography, behavioral assessment, and whole-mount imaging of diaphragmatic innervation. We aimed to determine whether ischemic stroke induces left–right diaphragmatic neuromuscular imbalance and to explore whether diaphragmatic electromyographic parameters are associated with motor recovery. We hypothesized that unilateral cerebral ischemia disrupts bilateral diaphragmatic activation, contributes to respiratory dysfunction, and reflects stroke-related neuromuscular impairment and recovery status.

## Methods

### Clinical Study Population and Data Collection

Clinical data were integrated from two independent cohorts of patients with mild acute ischemic stroke and healthy controls previously collected by our group using a standardized protocol. The study was registered at the Chinese Clinical Trial Registry (ChiCTR-2000031379). Recruitment at each participating center was initiated only after approval by the local institutional review board, and written informed consent was obtained from all participants.

Demographic characteristics, stroke-related variables, and baseline cardiorespiratory fitness indicators were independently abstracted by two researchers. To ensure data completeness and consistency, analyses were restricted to participants with complete records in both datasets. For cardiopulmonary exercise testing (CPET) variables, including peak oxygen uptake (VO₂peak), anaerobic threshold, myocardial rate-pressure product, peak circulatory power, and ventilatory dead space fraction, only participants who completed CPET and had complete records from both researchers were included. Participants unable to complete the maximal CPET protocol because of moderate-to-severe motor deficits, severe ataxia, cognitive impairment, or safety concerns were excluded. Additional details are provided in the Supplemental Methods.

### Animals and Experimental Stroke Model

Male C57BL/6J mice aged 6 to 8 weeks were used for animal experiments. All procedures were approved by the Animal Welfare and Ethics Committee of the Department of Laboratory Animal Science, Fudan University, and were performed in accordance with institutional guidelines for the care and use of laboratory animals.

Transient focal cerebral ischemia was induced by intraluminal filament occlusion of the left middle cerebral artery. Cerebral blood flow was monitored by laser speckle contrast imaging before occlusion, during ischemia, and after reperfusion. Mice with insufficient cerebral blood flow reduction during occlusion were excluded. Neurological deficits were assessed after stroke using the modified Neurological Severity Score. Detailed surgical procedures and exclusion criteria are described in the Supplemental Methods.

### Treadmill exercise test

Whole-body metabolic and cardiorespiratory responses to graded treadmill exercise were measured using a metabolic treadmill system. Oxygen consumption and carbon dioxide production were continuously recorded during incremental exercise, and VO₂peak was used as the primary index of exercise capacity. The treadmill protocol and exhaustion criteria are described in detail in the Supplemental Methods.

### Diaphragm Ultrasound, Diaphragmatic EMG, and Behavioral Assessments

Diaphragm excursion and thickness were assessed in vivo using high-frequency ultrasonography at predefined post-stroke time points. Bilateral diaphragmatic electromyography was recorded from the left and right hemidiaphragms with simultaneous electrocardiographic and respiratory monitoring. ECG artifacts were removed using an R-peak-locked template-subtraction approach, and dEMG peak amplitude, area under the curve, discharge duration, and EMG-respiration timing difference were quantified using custom MATLAB scripts. Motor function was evaluated using the modified Neurological Severity Score, rotarod test, and left and right forelimb grip strength test. Detailed procedures for ultrasound imaging, dEMG acquisition and processing, and behavioral assessments are provided in the Supplemental Methods.

### Whole-Mount Diaphragm Clearing and Light-Sheet Imaging

Whole-mount diaphragms were optically cleared, immunostained for the neuronal marker Tubb3/Tuj1, and imaged by light-sheet fluorescence microscopy. Tuj1-positive nerve fibers were visualized in three-dimensional z-stack images and quantified separately in the left and right hemidiaphragms. Detailed staining, clearing, imaging, and quantification procedures are provided in the Supplemental Methods.

### Statistical Analysis

Statistical analyses were performed using GraphPad Prism 10.1.2. Clinical continuous variables are presented as mean ± standard deviation, whereas animal experimental data are presented as mean ± standard error of the mean. Categorical variables are expressed as frequencies or percentages. Data normality was assessed using the Shapiro-Wilk test. Comparisons between two groups were performed using unpaired two-tailed Student’s t tests for normally distributed data or Mann-Whitney U tests for non-normally distributed data. Comparisons among multiple groups were performed using one-way or two-way ANOVA followed by Tukey’s multiple comparisons test, or nonparametric tests followed by Dunn’s multiple comparisons test, as appropriate. Repeated-measures or unbalanced longitudinal data were analyzed using mixed-effects models followed by Tukey’s multiple comparisons test. Correlations between behavioral, ultrasound, and dEMG parameters were assessed using Pearson correlation coefficients. A two-sided P value <0.05 was considered statistically significant.

## Results

### CPET identifies Cardiorespiratory impairment after stroke

To evaluate whether stroke is associated with measurable cardiorespiratory dysfunction, we analyzed CPET data from two previously established cohorts according to predefined data-completeness criteria. The final analysis included 22 patients with mild acute ischemic stroke and 15 age- and sex-matched healthy controls. Among the stroke group, 8 patients were female and 14 were male; the mean time since stroke onset was 108.8 ± 92.08 days; 9 patients had left-sided lesions and 13 had right-sided lesions. Detailed demographic and baseline characteristics are shown in Fig. 1A, Table 1, and Supplementary Fig. S1. Compared with healthy controls, patients with stroke exhibited significantly reduced peak oxygen uptake (VO₂ peak) and anaerobic threshold, indicating diminished exercise capacity. Stroke patients also showed an altered dead space-to-tidal volume ratio at peak exercise, suggesting impaired ventilatory efficiency. In addition, group differences were observed in mean rate-pressure product and pulmonary capillary pressure, reflecting abnormal cardiovascular and pulmonary circulatory responses during exercise (Fig. 1B–M). Together, these findings indicate that even mild ischemic stroke is associated with detectable impairment in exercise capacity, ventilatory efficiency, and cardiopulmonary regulation.

**Figure 1.**
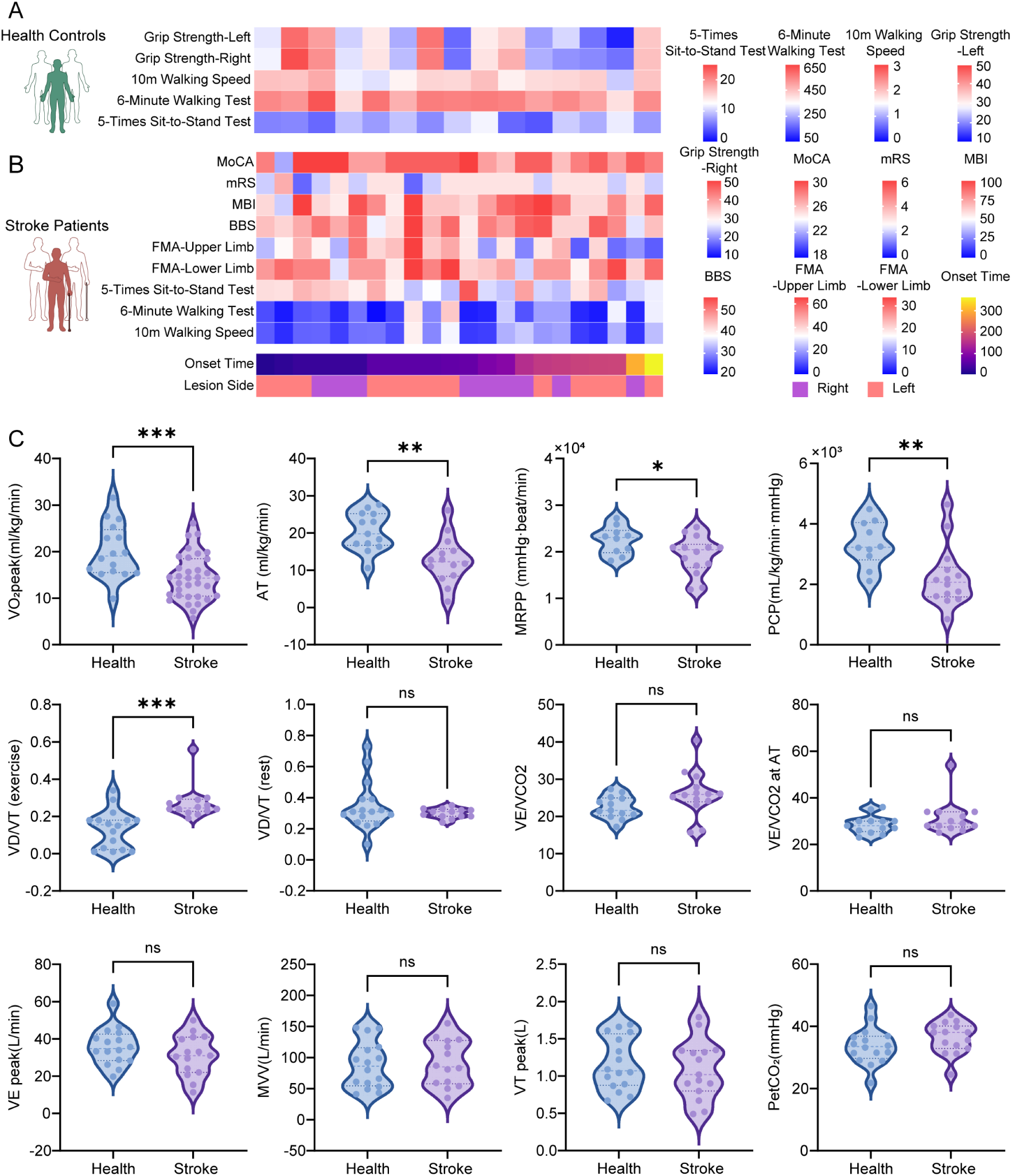
Clinical characteristics and CPET-derived cardiorespiratory dysfunction after stroke. (A-B) Heatmap showing individual-level clinical and functional characteristics of healthy controls(A) and patients with stroke(B), including grip strength, 10 m walking speed, 6 min walking test, 5-times sit-to-stand test, Montreal Cognitive Assessment, modified Rankin Scale, Modified Barthel Index, Berg Balance Scale, Fugl–Meyer Assessment scores, onset time, and lesion side. (C) Violin plots comparing cardiopulmonary exercise testing parameters between healthy controls and patients with stroke. Statistical analyses were performed using unpaired Student’s t-test for normally distributed data and Mann-Whitney U test for non-normally distributed data. Statistical significance is indicated as *P < 0.05, **P < 0.01, ***P < 0.001; ns, not significant. Abbreviations: VO₂peak, peak oxygen uptake normalized to body weight; AT, anaerobic threshold; MRPP, metabolic rate pressure product; PCP, peak circulatory power; VD/VT exercise, dead space to tidal volume ratio during exercise; VD/VT rest, dead space to tidal volume ratio at rest; VE/VCO₂, ventilatory equivalent for carbon dioxide; VE/VCO₂ at AT, VE/VCO₂ at anaerobic threshold; VE peak, peak minute ventilation; MVV, maximal voluntary ventilation; VT peak, peak tidal volume; PetCO₂, end-tidal carbon dioxide pressure.

**Table 1.** Demographic and clinical characteristics related to cardiorespiratory fitness (CRF) of the two groups. Continuous variables are presented as mean ± standard deviation, and categorical variables as counts. Differences between the two groups were analyzed using unpaired Student’s t-test for normally distributed data and Mann-Whitney U test for non-normally distributed data. Abbreviations: BMI, body mass index; FTSST, five-times sit-to-stand test; 6MWD, 6-min walk distance; 10mWT, 10-m walk test.

|  | Patients with stroke | Health subjects | P-value |
| --- | --- | --- | --- |
|  | N = 34 | N=16 |  |
| Sex (F/M) | 8/14 | 9/6 | 0.1931 |
| Age (years) | $53.5 \pm 14.06$ | $58.40 \pm 13.50$ | 0.2956 |
| BMI ($\text{kg}/\text{m}^2$) | $24.14 \pm 2.887$ | $23.53 \pm 3.072$ | 0.5426 |
| FTSST (s) | $13.51 \pm 3.892$ | $8.003 \pm 2.381$ | <0.0001 |
| 6MWD (m) | $184.6 \pm 109.6$ | $517.9 \pm 58.01$ | <0.0001 |
| 10mWT (m/s) | $0.7195 \pm 0.3915$ | $1.754 \pm 0.2767$ | <0.0001 |

### Diaphragmatic functional impairment and bilateral EMG imbalance after experimental stroke

Treadmill-based VO₂peak testing was performed to evaluate exercise capacity in mice after MCAO. VO₂ peak declined after MCAO, with the most pronounced reduction observed during the early post-stroke phase, followed by partial recovery over time (Fig. 2A-B). Mortality was also observed during the observation period, with the highest loss in the acute and subacute phases after cerebral ischemia (Fig. 2C). We next assessed diaphragmatic structure and mechanical function by ultrasonography. Diaphragmatic excursion amplitude was significantly altered at 24 h and 1 week after MCAO, whereas diaphragmatic thickness did not differ significantly at either time point (Fig. 2D). These findings indicate experimental stroke impaired diaphragmatic mechanics within the observed time window, without detectable diaphragmatic thinning. To determine whether diaphragmatic mechanical dysfunction was accompanied by altered neuromuscular activation, bilateral dEMG was recorded together with simultaneous respiratory flow monitoring after MCAO (Fig. 2E). EMG parameters, including peak amplitude, area under the curve (AUC), and discharge duration, were quantified at 24 h, 1 week, and 2 weeks after MCAO (Fig. 2F-H). MCAO induced side-dependent changes in diaphragmatic EMG activity. Across post-stroke time points, the left hemidiaphragm generally exhibited lower EMG activity than the contralateral right hemidiaphragm, yielding a relative right-dominant activation pattern that was most evident at 1 and 2 weeks after MCAO. Temporal analysis further showed that EMG parameters increased from 24 h to 1 week after ischemia and declined thereafter by 2 weeks. However, comparisons with baseline measurements or control of animals did not demonstrate a significant absolute increase in right hemi-diaphragmatic EMG activity (Fig. S2C-E), suggesting that the observed right-dominant pattern primarily reflected relative bilateral imbalance rather than overt contralateral hyperactivation. Together, these results demonstrate that left MCAO causes reduced diaphragmatic excursion and side-dependent diaphragmatic neuromuscular imbalance in the absence of detectable diaphragmatic thinning.

**Figure 2.**
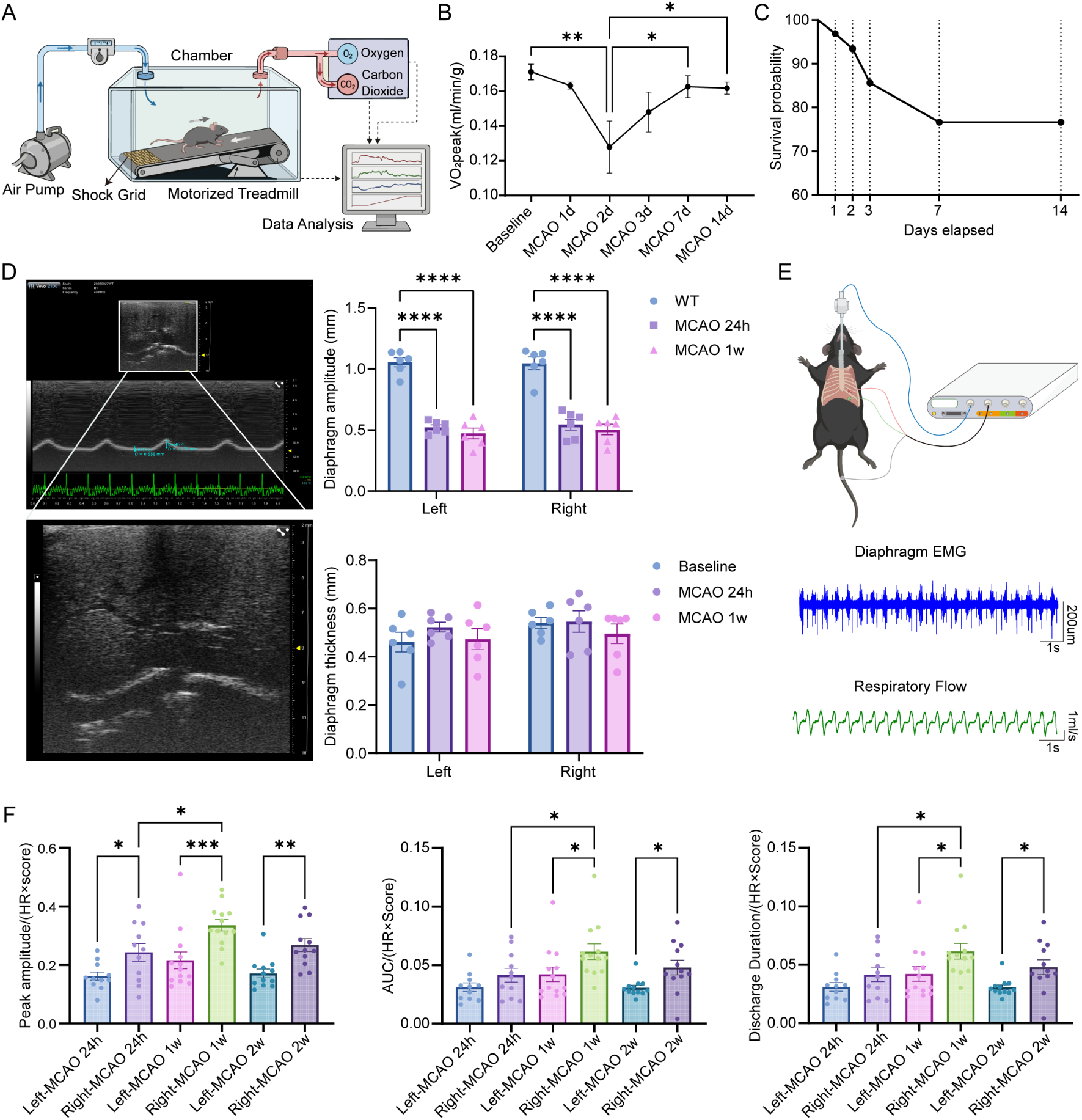
Post-stroke mice exhibit impaired exercise capacity, diaphragmatic dysfunction, and bilateral EMG imbalance. (A) Schematic illustration of the treadmill-based VO₂peak testing system. (B) Quantification of body weight-normalized VO₂peak at baseline (n = 10 mice) and 1 d (n = 4 mice), 2 d (n = 4 mice), 3 d (n = 5 mice), 7 d (n = 8 mice), and 14 d (n = 11 mice) after MCAO. (C) Kaplan–Meier survival curve of MCAO mice during the experimental period following treadmill-based maximal exercise testing. (D) Representative diaphragm ultrasound images and quantification of diaphragm excursion amplitude and diaphragm thickness in control and MCAO mice (n = 6 mice per group). (E) Schematic diagram of in situ dEMG recording with simultaneous respiratory flow monitoring, with representative EMG and respiratory flow traces. (F) Quantification of in situ dEMG parameters in the left and right hemidiaphragms at 24 h (n = 11 mice), 1 week (n = 13 mice), and 2 weeks (n = 12 mice) after MCAO, including peak amplitude normalized to HR × mNSS score, AUC of EMG discharge normalized to HR × mNSS score, and discharge duration normalized to HR × mNSS score. Data are presented as mean ± SEM with individual data points. Statistical analysis was performed using one-way ANOVA followed by Tukey’s multiple comparisons test for panel B, two-way ANOVA followed by Tukey’s multiple comparisons test for panel D, and Kruskal– Wallis test followed by Dunn’s multiple comparisons test for panel F. Statistical significance is indicated as *P < 0.05, **P < 0.01, ***P < 0.001, and ****P < 0.0001. Abbreviations: VO₂peak, peak oxygen uptake; AUC, area under the curve; HR, heart rate.

### Asymmetric remodeling of diaphragmatic innervation

To determine whether the post-stroke imbalance dEMG activity was accompanied by structural alterations in diaphragmatic innervation, we used whole-mount immunofluorescence staining for the pan-neuronal marker Tuj1 combined with tissue clearing to visualize the diaphragmatic intramuscular nerve network (Fig. 3A, B). Three-dimensional imaging revealed progressive remodeling of intramuscular innervation in both hemidiaphragms after MCAO. Quantitative analysis showed time-dependent changes in nerve fiber density with distinct trajectories between the two sides (Fig. 3C). At 24 h after MCAO, nerve fiber density was higher in the contralateral right hemidiaphragm than in the ipsilesional left hemidiaphragm. By 1 week, nerve fiber density in the right hemidiaphragm had markedly decreased, whereas the left hemidiaphragm was relatively preserved, thereby attenuating the initial side difference. At 2 weeks, nerve fiber density was reduced bilaterally, with no clear inter-hemidiaphragmatic difference remaining. These findings indicate that left MCAO is accompanied by time- and side-dependent remodeling of the diaphragmatic intramuscular nerve network.

**Figure 3.**
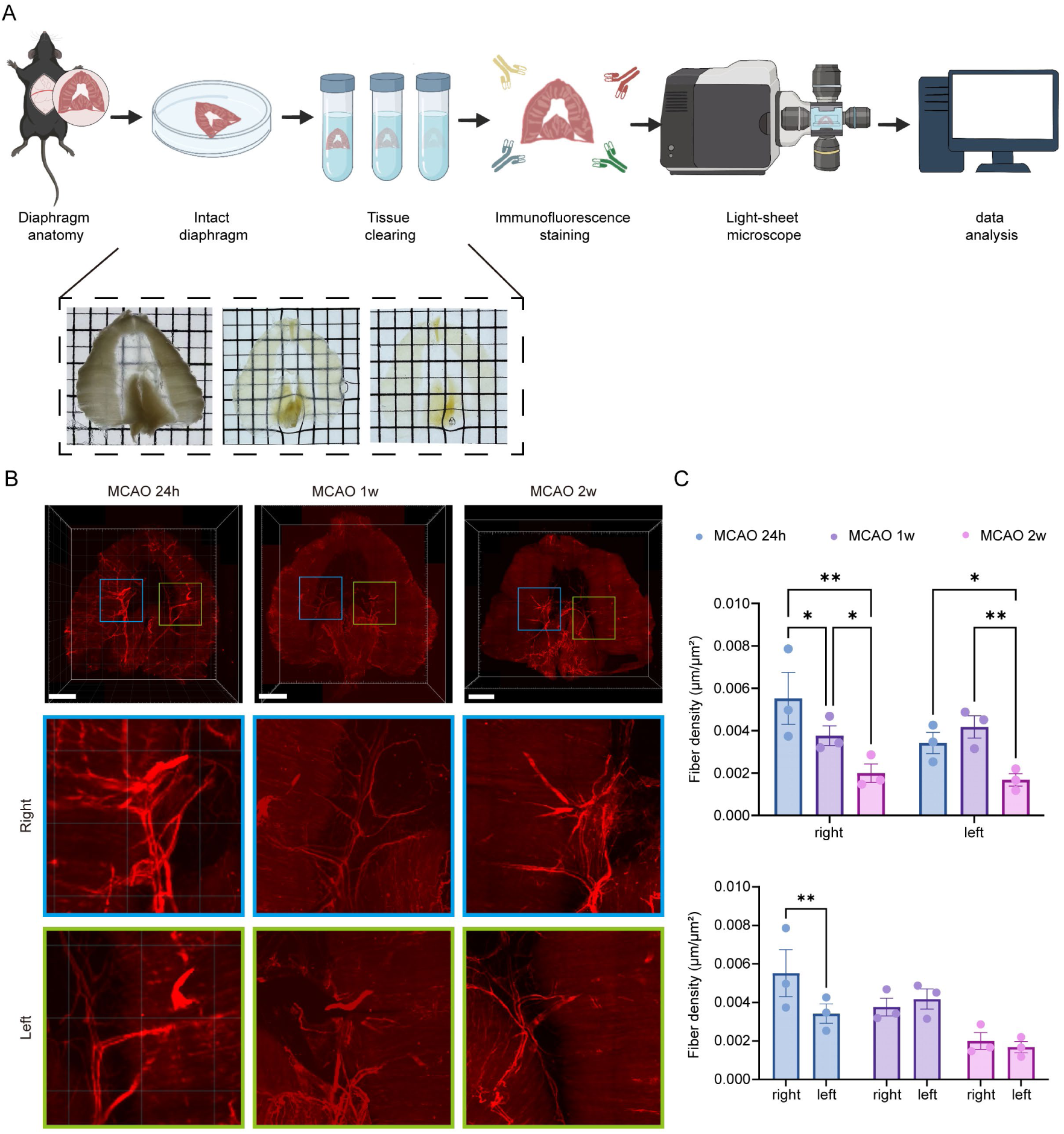
Asymmetric remodeling of diaphragmatic innervation after experimental stroke. (A) Schematic workflow of whole-mount diaphragm tissue clearing, Tuj1 immunofluorescence staining, light-sheet microscopy imaging, and quantitative image analysis. Representative images show the intact diaphragm before and after tissue clearing. (B) Representative light-sheet immunofluorescence images of whole-mount diaphragms stained with the neuronal marker Tuj1 at 24 h, 1 week, and 2 weeks after MCAO. Enlarged views of the right and left hemidiaphragms regions are shown below. Scale bar, 3000 µm. (C) Quantitative analysis of nerve fiber density in the right and left hemidiaphragms at 24 h, 1 week, and 2 weeks after MCAO (n = 3 mice per group). Data are presented as mean ± SEM with individual data points. Statistical analysis was performed using two-way ANOVA followed by Tukey’s multiple comparisons test for panel C. Statistical significance is indicated as *P < 0.05, **P < 0.01.

### Associations between diaphragmatic parameters and post-stroke functional recovery

To determine whether diaphragmatic functional indices reflect overall functional recovery after ischemic stroke, forelimb grip strength and rotarod performance were assessed serially following left MCAO (Fig. 4A–C). Forelimb grip strength declined markedly during the early post-stroke period, accompanied by clear interlimb functional asymmetry in the acute and subacute phases. Motor performance gradually improved over time, consistent with the spontaneous recovery profile typically observed in the MCAO model. We next examined the relationship between diaphragmatic ultrasound parameters and behavioral motor outcomes. Diaphragmatic excursion amplitude showed positive trends with bilateral forelimb grip strength and rotarod latency to fall, whereas diaphragmatic thickness showed negative trends with these motor measures (Fig. 4D). These findings suggest that impaired diaphragmatic mechanics may be associated with poorer functional performance after stroke.

**Figure 4.**
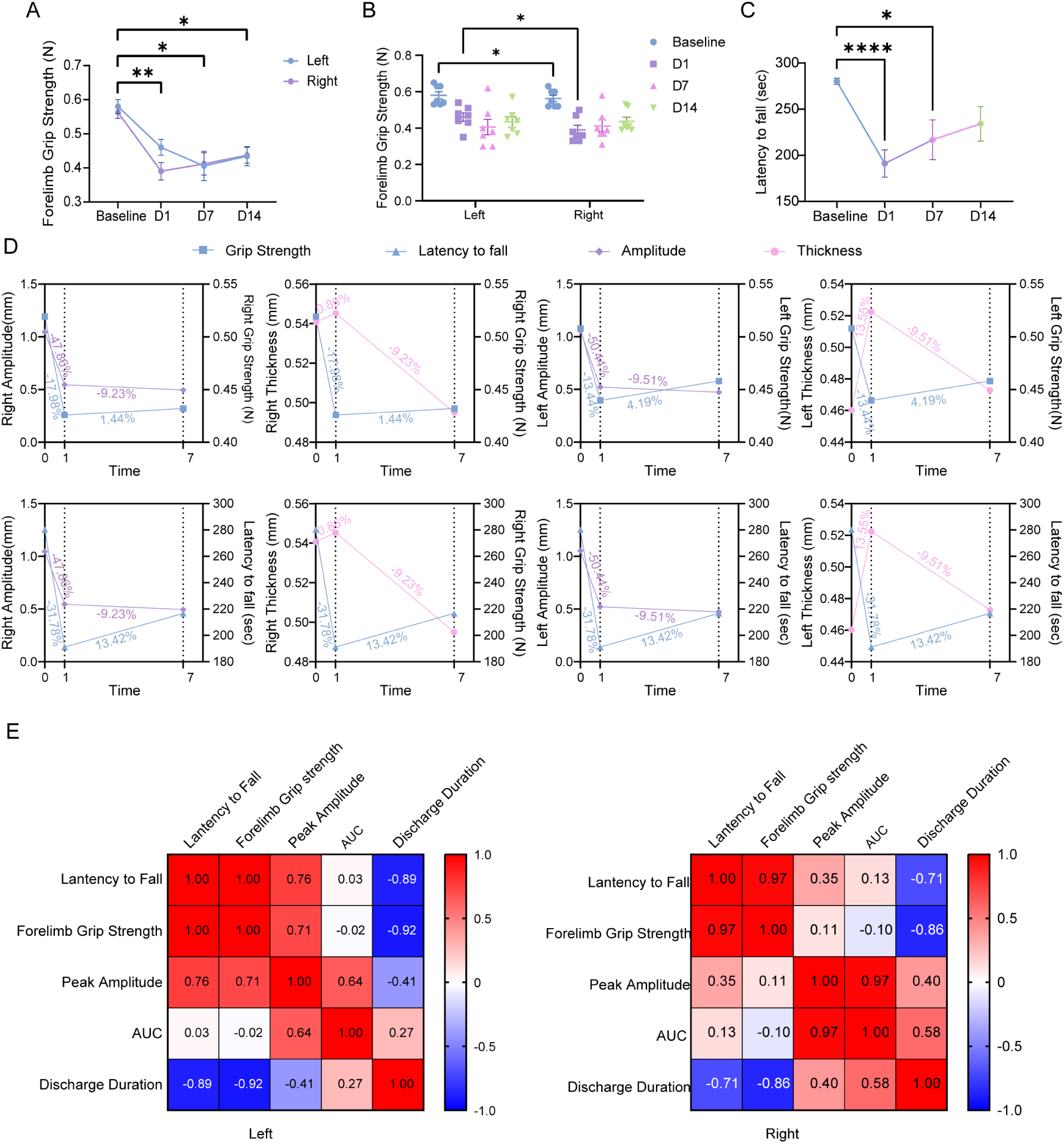
Associations between motor performance and diaphragm ultrasound or EMG parameters after MCAO in mice. (A) Changes in left and right forelimb grip strength at baseline and at 1, 7, and 14 days after MCAO (n = 7 mice). (B) Comparison of left and right forelimb grip strength at different time points after MCAO (n = 7 mice). (C) Rotarod performance assessed by latency to fall at baseline (n = 41 mice) and at 1 d (n = 41 mice), 7 d (n = 15 mice), and 14 d (n = 13 mice) after MCAO. (D) Temporal changes in diaphragm ultrasound parameters, including diaphragm excursion amplitude and diaphragm thickness, in relation to forelimb grip strength and latency to fall. Data are shown separately for the left and right hemidiaphragms(n = 4 mice). (E) Correlation matrices showing the relationships between motor performance and dEMG parameters in the left and right hemidiaphragms after MCAO(n = 4 mice). The color scale represents the strength and direction of the correlation, ranging from −1.0 to 1.0. Red indicates positive correlations, blue indicates negative correlations, and white indicates weak or no correlation. Data are presented as mean ± SEM with individual data points. Statistical analyses were performed using two-way ANOVA followed by Tukey’s multiple comparisons test for panels A and B, and a mixed-effects model followed by Tukey’s multiple comparisons test for panel C. Correlations in panels D and E were assessed using Pearson correlation coefficients. Statistical significance is indicated as *P < 0.05, **P < 0.01, and ****P < 0.0001.

To further assess the potential prognostic relevance of diaphragmatic electrophysiological activity, exploratory correlation heatmap analyses were performed to examine pairwise associations between dEMG parameters and post-stroke motor outcomes, separately for the left and right hemidiaphragms (Fig. 4E). None of the correlations reached statistical significance; therefore, all findings from this analysis should be interpreted as exploratory. In the ipsilesional left hemidiaphragm, peak amplitude showed positive trends with both rotarod latency to fall and forelimb grip strength, whereas AUC showed little apparent linear association with either outcome. By contrast, discharge duration showed negative trends with both behavioral measures. In the contralateral right hemidiaphragm, peak amplitude demonstrated weaker positive trends with motor performance, while discharge duration again showed negative trends with both outcomes. Overall, the correlation pattern suggested a tentative positive association between ipsilesional EMG amplitude and motor recovery, as well as an inverse association between discharge duration and behavioral performance across both sides.

## Discussion

In this study, complementary clinical and experimental data support the concept that respiratory dysfunction after stroke is not limited to reduced global exercise capacity but also involves altered diaphragmatic motor control. Clinically, CPET showed that even patients with mild ischemic stroke had impaired exercise capacity and abnormal cardiopulmonary responses. Experimentally, unilateral cerebral ischemia in mice was associated with reduced diaphragmatic excursion, bilateral asymmetry in diaphragmatic EMG activity, and time-dependent remodeling of the diaphragmatic intramuscular nerve network. Taken together, these findings suggest that post-stroke respiratory dysfunction may include a lateralized disturbance of inspiratory neuromuscular output in addition to global cardiorespiratory impairment.

A notable finding was that the diaphragmatic EMG phenotype after left MCAO was characterized by a relative right-dominant activation pattern that appeared to arise primarily from reduced ipsilesional activity rather than clear contralesional hyperactivation. This distinction is mechanistically relevant. A true increase in contralesional activity would favor a compensatory recruitment model, whereas the present pattern is more consistent with impaired or dysregulated ipsilesional descending drive leading to interhemidiaphragmatic imbalance. This interpretation aligns with prior clinical observations showing that stroke-related diaphragmatic dysfunction is often more prominent on the hemiparetic or lesion-contralateral side, although bilateral abnormalities may also occur depending on lesion location and respiratory task demands ^13,18^.

The ultrasound findings further support a predominantly functional, rather than structural, abnormality in the early phase after stroke. Diaphragmatic excursions were reduced, whereas diaphragmatic thickness was preserved. Reduced excursion in the absence of detectable thinning is more consistent with impaired neural activation or contractile recruitment than with early muscle atrophy^22^. This interpretation is in line with previous stroke studies and recent reviews indicating that dynamic ultrasound indices, including excursion and thickening fraction, are more sensitive to early diaphragmatic dysfunction than static thickness measurements alone ^11,23–25^. The present data therefore supports the view that abnormalities in diaphragmatic motor drive may emerge before overt structural loss of muscle mass.

The neurophysiological findings are biologically plausible in the context of the complex central control of the diaphragm. Unlike limb muscles, diaphragmatic activity is generated through the interaction of automatic bulbospinal respiratory circuits and supraspinal pathways involved in volitional breathing, postural control, and respiratory-motor integration^17^. Previous studies using magnetic stimulation have demonstrated abnormal corticodiaphragmatic conduction and reduced diaphragmatic movement in acute ischemic stroke, and more recent work has confirmed the feasibility of assessing corticodiaphragmatic output using motor evoked responses ^15^. Within this framework, unilateral MCAO may disrupt bilateral phrenic motor output through altered supraspinal modulation, even if the resulting pattern does not mirror the simple contralateral organization typically seen in limb motor deficits^26^.

The whole-mount imaging findings extend these physiological observations by showing that post-stroke diaphragmatic imbalance is accompanied by remodeling of the intramuscular nerve network. Importantly, these structural changes should not be interpreted as a direct surrogate for instantaneous EMG activity. Diaphragmatic EMG captures dynamic neural activation during breathing, whereas cleared whole-mount imaging reflects the anatomical state of intramuscular innervation over a longer timescale^27^. The divergence in temporal resolution between these measures may therefore be informative. One possible interpretation is that unilateral cerebral ischemia first perturbs bilateral respiratory motor drive, after which side- and time-dependent peripheral neural remodeling develops secondarily^28^. This interpretation is compatible with broader evidence that respiratory muscle properties are shaped by neural activity patterns and that the diaphragm functions within an integrated sensorimotor network rather than as an isolated ventilatory pump^16^.

At a systems level, the present findings may also reflect remote network effects of focal cerebral injury. Stroke can induce diaschisis and alter interhemispheric interactions, thereby disrupting function in anatomically intact but connected regions ^20,29–31^. Although such mechanisms are most often discussed in the context of limb recovery or cognition, they are equally relevant to respiratory control, which depends on coordinated interactions among cortical, subcortical, brainstem, and spinal circuits^32^. The observed diaphragmatic asymmetry may therefore represent a network-level consequence of unilateral stroke rather than a purely local lesion effect.

The exploratory association analyses between diaphragmatic measures and motor behavior should be interpreted cautiously. None of the correlations between dEMG parameters and behavioral outcomes reached statistical significance, and these analyses were therefore hypothesis-generating. Nevertheless, the directional pattern was of interest: higher ipsilesional EMG amplitude tended to be associated with better motor performance, whereas longer discharge duration tended to be associated with poorer performance on both sides. Although preliminary, these findings raise the possibility that more efficient inspiratory recruitment may parallel with broader recovery of sensorimotor function. This possibility is supported indirectly by clinical studies linking diaphragmatic or respiratory muscle measures to limb motor performance, balance, and activities of daily living after stroke^21^. Whether EMG-derived diaphragmatic indices have prognostic value will require validation in larger longitudinal datasets.

These findings also help distinguish the potential roles of CPET and diaphragm-focused physiological assessment. CPET provides an integrated measure of cardiorespiratory impairment and demonstrates clinically relevant abnormalities even in mild stroke^33,34^. However, CPET requires cooperation, preserved mobility, and sufficient exercise tolerance. By contrast, diaphragm ultrasound and dEMG assess respiratory motor function more directly and may remain feasible when maximal exercise testing is impractical^35^. This distinction is clinically relevant given growing evidence from systematic reviews and meta-analyses that respiratory muscle training and aerobic exercise can improve function after stroke^5,9,34^. Earlier identification of specific respiratory motor abnormalities may therefore help refine rehabilitation strategies.

Several limitations warrant consideration. First, the link between cortical ischemic injury and altered phrenic motor output was inferred rather than directly demonstrated. Future studies should combine lesion characterization with cortico-diaphragmatic stimulation, phrenic electrophysiology, and neuromuscular junction analyses. Second, Tuj1 staining visualizes the intramuscular neural network but does not distinguish motor from sensory or autonomic fibers, nor does it define synaptic integrity at the neuromuscular junction. Third, the behavioral correlation analyses were exploratory and likely underpowered. Fourth, absolute dEMG values may have been influenced by anesthesia, surgical preparation, and signal normalization procedures. These methodological factors should be further optimized in future work.

In conclusion, the present study supports a model in which post-stroke diaphragmatic dysfunction includes a lateralized neuromuscular imbalance in addition to reduced overall cardiorespiratory performance. Bilateral diaphragmatic EMG provides a functional readout of disrupted left-right inspiratory activation in experimental stroke, and whole-mount nerve imaging suggests that this imbalance is accompanied by dynamic remodeling of diaphragmatic intramuscular innervation. These findings broaden the current understanding of respiratory dysfunction after stroke and support further investigation of diaphragmatic electrophysiological measures as markers of post-stroke respiratory motor control and recovery-related neuromuscular integrity.

## References

1. Collaborators GBDSRF. Global, regional, and national burden of stroke and its risk factors, 1990-2021: a systematic analysis for the Global Burden of Disease Study 2021. Lancet Neurol. 2024;23:973–1003. doi: 10.1016/S1474-4422(24)00369-7

2. Rochester CL, Vogiatzis I, Holland AE, Lareau SC, Marciniuk DD, Puhan MA, Spruit MA, Masefield S, Casaburi R, Clini EM, et al. An Official American Thoracic Society/European Respiratory Society Policy Statement: Enhancing Implementation, Use, and Delivery of Pulmonary Rehabilitation. Am J Resp Crit Care. 2015;192:1373–1386. doi: 10.1164/rccm.201510-1966ST

3. Fan QW, Jia J. Translating Research Into Clinical Practice Importance of Improving Cardiorespiratory Fitness in Stroke Population. Stroke. 2020;51:361–367. doi: 10.1161/Strokeaha.119.027345

4. Abdullahi A, Wong TWL, Ng SSM. Efficacy of diaphragmatic breathing exercise on respiratory, cognitive, and motor function outcomes in patients with stroke: a systematic review and meta-analysis. Frontiers in Neurology. 2024;14:1233408. doi: 10.3389/fneur.2023.1233408

5. Fabero-Garrido R, Del Corral T, Angulo-Diaz-Parreno S, Plaza-Manzano G, Martin-Casas P, Cleland JA, Fernandez-de-Las-Penas C, Lopez-de-Uralde-Villanueva I. Respiratory muscle training improves exercise tolerance and respiratory muscle function/structure post-stroke at short term: A systematic review and meta-analysis. Ann Phys Rehabil Med. 2022;65:101596. doi: 10.1016/j.rehab.2021.101596

6. Huang L, Zhang JM, Bi ZT. Effects of respiratory muscle training on respiratory function, exercise capacity, and quality of life in chronic stroke patients: a systematic review and meta-analysis. Frontiers in Physiology. 2025;16:1642262. doi: 10.3389/fphys.2025.1642262

7. Billinger SA, Arena R, Bernhardt J, Eng JJ, Franklin BA, Johnson CM, MacKay-Lyons M, Macko RF, Mead GE, Roth EJ, et al. Physical Activity and Exercise Recommendations for Stroke Survivors A Statement for Healthcare Professionals From the American Heart Association/American Stroke Association. Stroke. 2014;45:2532–2553. doi: 10.1161/Str.0000000000000022

8. Saunders DH, Sanderson M, Hayes S, Johnson L, Kramer S, Carter D, Jarvis H, Brazzelli M, Mead GE. Physical Fitness Training for Patients With Stroke. Stroke. 2020;51:E299–E300. doi: 10.1161/Strokeaha.120.030826

9. Zhang W, Pan H, Zong Y, Wang J, Xie Q. Respiratory Muscle Training Reduces Respiratory Complications and Improves Swallowing Function After Stroke: A Systematic Review and Meta-Analysis. Arch Phys Med Rehabil. 2022;103:1179– 1191. doi: 10.1016/j.apmr.2021.10.020

10. Machado N, Williams G, Olver J, Johnson L. The safety and feasibility of early cardiorespiratory fitness testing after stroke. Pm&R. 2023;15:291–301. doi: 10.1002/pmrj.12787

11. Liu X, Yang Y, Jia J. Respiratory muscle ultrasonography evaluation and its clinical application in stroke patients: a review. Front Neurosci-Switz. 2023;17:1132335. doi: 10.3389/fnins.2023.1132335

12. Del Negro CA, Funk GD, Feldman JL. Breathing matters. Nat Rev Neurosci. 2018;19:351–367. doi: 10.1038/s41583-018-0003-6

13. Catala-Ripoll JV, Monsalve-Naharro JA, Hernandez-Fernandez F. Incidence and predictive factors of diaphragmatic dysfunction in acute stroke. BMC Neurology. 2020;20:79. doi: 10.1186/s12883-020-01664-w

14. Khedr EM, El Shinawy O, Khedr T, Aziz Ali YA, Awad EM. Assessment of corticodiaphragmatic pathway and pulmonary function in acute ischemic stroke patients. European Journal of Neurology. 2000;7:323–330. doi: 10.1046/j.1468-1331.2000.00078.x

15. Welch JF, Argento PJ, Mitchell GS, Fox EJ. Reliability of diaphragmatic motor-evoked potentials induced by transcranial magnetic stimulation. Journal of Applied Physiology. 2020;129:1393–1404. doi: 10.1152/japplphysiol.00486.2020

16. Kocjan J, Adamek M, Gzik-Zroska B, Czyzewski D, Rydel M. Network of breathing. Multifunctional role of the diaphragm: a review. Advances in Respiratory Medicine. 2017;85:224–232. doi: 10.5603/ARM.2017.0037

17. Fuller DD, Rana S, Smuder AJ, Dale EA. Chapter 15 - The phrenic neuromuscular system. In: Chen R, Guyenet PG, eds. Handbook of Clinical Neurology. Elsevier; 2022:393–408.

18. Liu F, Jones AYM, Tsang RCC, Yam TTT, Tsang WWN. Diaphragm and sternocleidomastoid muscle activity with increasing inspiratory pressure loads in people after stroke. Scientific Reports. 2025;15:5856. doi: 10.1038/s41598-025-90199-6

19. Liu X, Qu Q, Deng P, Zhao Y, Liu C, Fu C, Jia J. Assessment of Diaphragm in Hemiplegic Patients after Stroke with Ultrasound and Its Correlation of Extremity Motor and Balance Function. In: Brain Sci. Switzerland; 2022.

20. Bundy DT, Nudo RJ. Preclinical Studies of Neuroplasticity Following Experimental Brain Injury An Update. Stroke. 2019;50:2626–2633. doi: 10.1161/Strokeaha.119.023550

21. Li M, Huang Y, Chen H. Relationship between motor dysfunction, the respiratory muscles and pulmonary function in stroke patients with hemiplegia: a retrospective study. BMC Geriatrics. 2024;24:59. doi: 10.1186/s12877-023-04647-x

22. Qi H, Tian D, Luan F, Yang R, Zeng N. Pathophysiological changes of muscle after ischemic stroke: a secondary consequence of stroke injury. Neural Regen Res. 2024;19:737–746. doi: 10.4103/1673-5374.382221

23. Chen Y. Effect of respiratory muscle training on diaphragm function in stroke patients: a systematic review and meta-analysis. Frontiers in Medicine. 2025;12:1694356. doi: 10.3389/fmed.2025.1694356

24. Liu F, Jones AYM, Tsang RCC, Yam TTT, Tsang WWN. Effects of inspiratory muscle training on respiratory function, diaphragmatic thickness, balance control, exercise capacity and quality of life in people after stroke: A randomized controlled trial protocol. In: PLoS One. United States; 2025:e0319899.

25. Truong D, Abo S, Whish-Wilson GA, D’Souza AN, Beach LJ, Mathur S, Mayer KP, Ntoumenopoulos G, Baldwin C, El-Ansary D, et al. Methodological and Clinimetric Evaluation of Inspiratory Respiratory Muscle Ultrasound in the Critical Care Setting: A Systematic Review and Meta-Analysis. Critical Care Medicine. 2023;51:E24–E36. doi: 10.1097/Ccm.0000000000005739

26. Goshgarian HG. Plasticity in respiratory motor control - Invited review: The crossed phrenic phenomenon: a model for plasticity in the respiratory pathways following spinal cord injury. Journal of Applied Physiology. 2003;94:795–810. doi: 10.1152/japplphysiol.00847.2002

27. Hérent C, Diem S, Fortin G, Bouvier J. Absent phasing of respiratory and locomotor rhythms in running mice. Elife. 2020;9. doi: ARTN e61919 10.7554/eLife.61919

28. Fogarty MJ, Mantilla CB, Sieck GC. Breathing: Motor Control of Diaphragm Muscle. Physiology. 2018;33:113–126. doi: 10.1152/physiol.00002.2018

29. Gerges ANH, Hordacre B, Di Pietro F, Moseley GL, Berryman C. Do adults with stroke have altered interhemispheric inhibition? A systematic review with meta-analysis. Journal of Stroke and Cerebrovascular Diseases. 2022;31:106494. doi: 10.1016/j.jstrokecerebrovasdis.2022.106494

30. Pascoa dos Santos F, Verschure PFMJ. Excitatory-inhibitory homeostasis and diaschisis: tying the local and global scales in the post-stroke cortex. Frontiers in Systems Neuroscience. 2022;15:806544. doi: 10.3389/fnsys.2021.806544

31. Jia Q, Sheng NN, Naeije G. Prevalence and Imaging Correlates of Cerebral Diaschisis After Ischemic Stroke: A Systematic Review and Meta-Analysis. Brain Sciences. 2025;16. doi: ARTN 50 10.3390/brainsci16010050

32. Benarroch EE. Brainstem integration of arousal, sleep cardiovascular, and respiratory control. Neurology. 2018;91:958–966. doi: 10.1212/Wnl.0000000000006537

33. Li DX, Zhou MC, Zha FB, Long JJ, Wang YL. Cardiorespiratory fitness in stroke patients with different activities of daily living levels: a cross-sectional study. Scientific Reports. 2025;15. doi: ARTN 14109 10.1038/s41598-025-97293-9

34. Moncion K, Rodrigues L, de las Heras B, Noguchi KS, Wiley E, Eng JJ, MacKay-Lyons M, Sweet SN, Thiel A, Fung J, et al. Cardiorespiratory Fitness Benefits of High-Intensity Interval Training After Stroke: A Randomized Controlled Trial. Stroke. 2024;55:2202–2211. doi: 10.1161/Strokeaha.124.046564

35. Haaksma ME, Smit JM, La Heldeweg M, Nooitgedacht JS, Atmowihardjo LN, Jonkman AH, de Vries HJ, Lim EHT, Steenvoorden T, Lust E, et al. Holistic Ultrasound to Predict Extubation Failure in Clinical Practice. Resp Care. 2021;66:994–1003. doi: 10.4187/respcare.08679

